# PIKfyve is an essential component of the endolysosomal pathway within photoreceptors and the retinal pigment epithelium

**DOI:** 10.1101/2025.09.15.676399

**Authors:** Karen Attia, Ifrah Anjum, Susanne Lingrell, Matthew D Benson, Ian M MacDonald, Jennifer C Hocking

## Abstract

Phosphoinositides (PIs) are a family of seven low abundance membrane lipids, each with distinct signaling functions. The phosphoinositide kinase PIKfyve generates phosphoinositide-3,5-bisphosphate (PI(3,5)P₂) and PI5P. Emerging evidence implicates PIKfyve in key cellular processes, including autophagy, phagocytosis, endosomal trafficking, lysosomal maintenance, and melanosome formation. Complete loss of PIKfyve function is embryonic lethal in model organisms. In humans, heterozygous mutations in *PIKFYVE* are associated with Fleck corneal dystrophy and congenital cataracts. In this study, we investigate the role of PIKfyve in photoreceptors and the adjacent retinal pigment epithelium (RPE), host to dynamic endolysosomal pathways required for enduring the high oxidative stress environment, transporting metabolites and phototransduction components, and the breakdown of outer segment discs. To assess PIKfyve function in the retina and RPE in our zebrafish model, we employed CRISPR/Cas9-mediated gene editing and pharmacological inhibition using the specific PIKfyve inhibitor apilimod. Loss of PIKfyve activity leads to RPE expansion characterized by the accumulation of LC3- and LAMP1-positive vacuoles, along with defects in phagosome degradation and minor changes to melanosome biogenesis. Photoreceptors deprived of PIKfyve function develop a single large vacuole in the inner segment, while the OS remains largely intact over the timespan analyzed. Electroretinography (ERG) recordings revealed complete visual impairment in *pikfyve* crispant larvae, and significantly reduced visual function in larvae treated with apilimod post embryogenesis. These findings highlight the critical role of PIKfyve in the development and homeostasis of the RPE and retina.

## 1. INTRODUCTION

Vision begins with the detection of light by retinal rod and cone photoreceptors, whose function and survival are dependent on the adjacent retinal pigment epithelium (RPE) (Goldberg et al., 2016; Lakkaraju et al., 2020; Yang et al., 2021). The RPE and retina are long-lived tissues that cannot regenerate in humans; however, the large light-capturing photoreceptor outer segment (OS) undergoes constant renewal through growth at the base matched by trimming at the tip (LaVail, 1976; Megaw, 2025; Young, 1967; Young & Bok, 1969). RPE cells engulf and digest the OS tips, returning components to the photoreceptors. While typically considered a form of phagocytosis, the term trogocytosis has been suggested as more appropriate given that the entire cell is not being engulfed (Paniagua et al., 2025; Umapathy et al., 2023). The RPE also forms the outer blood-retinal barrier, transports ions and metabolites, recycles visual pigment, contains melanosomes to absorb stray light, and secretes growth factors and cytokines. RPE dysfunction and degeneration leads to loss of photoreceptors and eventual blindness (Gu et al., 2012; Keeling et al., 2018; Kiser & Palczewski, 2021).

Critical to RPE health is the endolysosomal pathway: the formation and resolution of endosomes, phagosomes, and autophagosomes (Etchegaray & Ravichandran, 2025; Intartaglia et al., 2022; Lakkaraju et al., 2020). Demand on the RPE endolysosomal pathway is enormous owing to the daily engulfment of POS tips as well as the endocytosis required for the transport of visual cycle components and metabolites between the RPE and retina (J.-Y. Kim et al., 2013; Saari, 2016; Sato & Kefalov, 2024; Wright et al., 2015). Further, the RPE and photoreceptors are among the most metabolically active cells in the body, which, combined with light exposure and the rapid perfusion rate of the adjacent choroidal vasculature, creates a high oxidative stress environment (Böhm et al., 2023; Hanna et al., 2022). Both cell types are therefore dependent on an active autophagy system for the frequent turnover of cellular organelles (Falcão et al., 2025; Intartaglia et al., 2022; Liton et al., 2023; Markitantova & Simirskii, 2025). Notably, lysosomal storage disorders are associated with progressive retinal degeneration (Boya et al., 2023).

Phosphoinositides are a class of low-abundance signaling lipids (Balla, 2013; Lolicato et al., 2024; Martin, 1998; Schink et al., 2016). The seven different subtypes each occupy specific cellular membranes, where they regulate protein localization and activity. The FYVE (Fab1, YOTB, Vac1, EEA1) zinc finger-containing phosphoinositide kinase PIKfyve phosphorylates the membrane lipid phosphatidylinositol 3-phosphate (PI3P), present on intracellular vesicles, into low-abundance PI(3,5)P2 (Fig. 1A) (Rivero-Ríos & Weisman, 2022; Zolov et al., 2012). PIKfyve can also generate PI5P, either directly by phosphorylation of PI or indirectly by providing PI(3,5)P2 for dephosphorylation, although the cellular functions of PI5P are less studied (Hasegawa et al., 2017; Rameh & Blind, 2023). PIKfyve is implicated in maturation and resolution of endosomes, autophagosomes, and phagosomes, and in the homeostasis and fission of lysosomes (Buckley et al., 2019; Dayam et al., 2017; G. H. E. Kim et al., 2014; Krishna et al., 2016; Rodgers et al., 2023; Takatori et al., 2016). A consistent phenotype upon PIKfyve inhibition is the buildup of swollen vacuoles.

**Figure 1.**
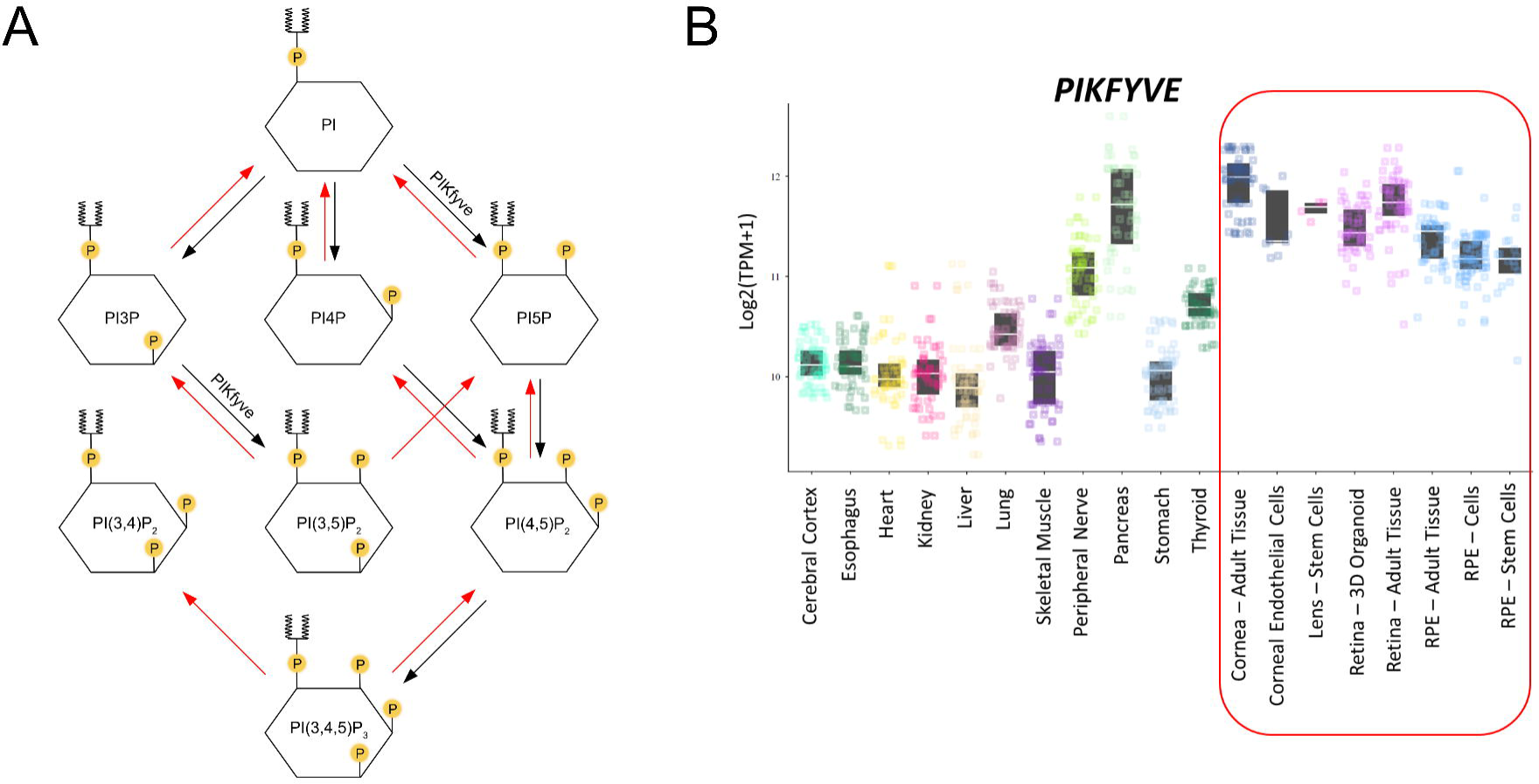
The phosphoinositide kinase *PIKFYVE* is expressed in human ocular tissues. A) The family of phosphoinositide lipids, highlighting the conversions mediated by PIKfyve to generate PI(3,5)P2 and PI(5)P. B) Expression of PIKfyve across select human tissue types based on RNASeq data collated in eyeIntegration v2.12, NEI, NIH. Note the high expression in ocular tissues (red box).

*PIKfyve* homozygous mutations are embryonic lethal in all model organisms tested, complicating analysis of function *in vivo* (Ikonomov et al., 2011; Lenk & Meisler, 2014; Mei et al., 2022). In humans, heterozygous mutations cause Fleck corneal dystrophy and are linked to congenital cataracts (Gee et al., 2015; Mei et al., 2022; Rodríguez-Solana et al., 2023). The role of PIKfyve in the RPE and retina has never been investigated, despite *PIKFYVE* transcript levels in the eye being among the highest of all human body tissues (Fig. 1B) and the known importance of the endolysosomal system for RPE function and overall eye health. Indeed, *PIKFYVE* was identified as a gene associated with age-related macular degeneration (AMD), the leading cause of vision loss in developed countries (Garcia-Garcia et al., 2022; Ulańczyk et al., 2021; Wąsowska et al., 2022). Of note, the PIKfyve inhibitor apilimod is being considered as a potential treatment for a variety of non-ocular diseases.

Here, we have investigated the consequences of PIKfyve loss in the zebrafish retina using two paradigms: *pikfyve* CRISPR injection into single-cell embryos (*pikfyve* crispants) and treatment of larvae with apilimod. We observed profound changes in the RPE of fish upon PIKfyve disruption, including the accumulation of vacuoles, impaired phagosome degradation, disrupted melanosome biogenesis, and expansion of the RPE impinging upon the photoreceptor layer. Photoreceptors were also affected, developing large inner segment vacuoles but generally maintaining normal OS layering. Finally, electroretinographic recordings revealed significant visual deficits upon PIKfyve loss. Our data highlights the importance of PIKfyve in the retina, with implications for disease risk and providing a consideration for the use of PIKfyve inhibitors as pharmaceuticals.

## 2. MATERIALS AND METHODS

### 2.1 Animal Ethics

Approval for this study was obtained from the University of Alberta Animal Care and Use Committee: Biosciences, under protocol AUP#1476.

### 2.2 Zebrafish Care

Zebrafish were maintained in a recirculating Tecniplast aquatics system and cared for by Health Sciences Laboratory Animal Services (HSLAS) at the University of Alberta. All solutions for live fish were made with reverse osmosis purified water. Stock embryonic media (EM) was prepared by combining 17.5 g NaCl, 0.75 g KCl, 2.9 g CaCl_2_-2H_2_O, 0.41 g KH_2_PO_4_, 0.142 g Na_2_HPO_4_ anhydrous, 4.9 g MgSO_4_-7H_2_O in 1 liter of water, followed by vacuum filtering and storage at 4°C. Working EM solution was prepared fresh every week by combining 50 mL stock EM, 2 mL sodium bicarbonate solution (0.3 g in 10 mL), and water to 1 liter, and stored at room temperature. Zebrafish embryos and larvae were grown in a 28.5°C incubator in a working EM solution up to 7 days post fertilization (dpf). Fish to be grown longer were transferred to the aquatics facility and fed three times per day as larvae and juveniles and twice per day as adults with a combination of rotifers and Ziegler dry food. Zebrafish were kept in a 14:10 hour light/dark cycle. Zebrafish lacking pigment in the eye and the body were generated by Spencer Balay in the laboratory of Dr. W. Ted Allison (University of Alberta) based on the previously described *crystal* line (Antinucci & Hindges, 2016; Balay, 2018). Tricaine mesylate (TMS) was used to anesthetize and euthanize zebrafish. A stock TMS solution was prepared by combining 400 g tricaine powder (3-aminobenzoic acid ethyl ester, MS222), 97.9 mL water, 2.1 mL 1M Tris pH 8 and stored at −20°C. A 4.2% working TMS solution for zebrafish anesthesia was made by dilution of the stock solution with EM. For euthanasia, anaesthetized fish were subsequently frozen or preserved in fixative.

### 2.3 CRISPR/Cas9 Mutagenesis

The CHOPCHOP web tool (https://chopchop.cbu.uib.no) and the Integrated DNA Technologies (IDT) Custom Alt-R CRISPR-Cas9 guide RNA tool were used for CRISPR RNA (crRNA) design. Two crRNAs were designed to target the third and fourth exons of the kinase domain of the *Danio rerio pikfyve* gene (Table S1). The first crRNA targets the sense strand of exon 38 while the second crRNA targets the antisense strand of exon 39 of transcript ENSDART00000184170.1. Co-injection of the two crRNAs potentially generates two double stranded breaks that are ∼268 nucleotides apart. crRNAs previously designed to target the pigment gene *slc45a2* were used as controls (Rosello et al., 2022) (Table S1). Alt-R ® CRISPR-Cas9 crRNAs, Alt-R ® CRISPR-Cas9 tracrRNA, and Alt-RTM S.p. Cas9 Nuclease V3 were ordered from IDT. RNAs were resuspended in RNase-free TE buffer (10 mM Tris, 0.1 mM EDTA) to a concentration of 100 μM. To anneal, 5 μL tracrRNA and 5 μL crRNA (100 μM each) were heated at 95°C for 5 minutes and then gradually cooled to room temperature in a thermocycler at 0.1°C/second. Annealed RNA duplexes were stored at −20°C. Injection mixtures consisted of 5 μM each RNA duplex diluted in IDT duplex buffer (30 mM HEPES, pH 7.5; 100 mM potassium acetate) and 5 μM Cas9 (diluted in 1X RNase-free PBS). Zebrafish embryos were injected at the single-cell stage with 2 nL of the injection mixture. To prepare injection needles, filamented borosilicate glass capillary tubes (OD 1.20mm, ID 0.90 mm; Sutter Instruments, Novato, USA) were pulled on a Sutter micropipette puller. Injections were performed under an Olympus stereo microscope using a micromanipulator and pneumatic pump (World Precision Instruments, Sarasota, USA). Injected embryos and uninjected siblings were grown in a 28.5°C incubator in EM for 1-7 dpf prior to genotyping and analysis or transfer to the fish facility.

### 2.4 Genotyping by Polymerase Chain Reaction

DNA was extracted from either individual embryos, pools of five to ten embryos, or adult fin clips by heating the tissue in 10 uL, 50 μL, or 100 uL 50 mM NaOH, respectively, at 95°C for 20 minutes before cooling to 4°C for 10 minutes and neutralizing with 1/10^th^ volume of 1M Tris-HCl, pH 8 (Meeker et al., 2007). PCR reactions were prepared by combining 10 μL 5X Green GoTaq Buffer, 1 μL nucleotide mix (1 mM each), 4 μL each of forward and reverse primer (5 μM each), 0.25 μL GoTaq Polymerase (Promega), <0.5 μg DNA, and water to 50 μL. The primers used to amplify the target region in PIKfyve were forward primer CAGACAGGGAACCCACATATT and reverse primer ACAGCCCTAGTTGGGACTAA. The PCR amplification reaction was set to 95°C for 2 minutes, 30 cycles of amplification (95°C for 15 seconds, 55°C for 30 seconds, 72°C for 15 seconds), 72°C for 4 minutes. PCR products were run on a 1% TAE buffer gel stained with Invitrogen 10,000X SYBR Safe DNA Gel Stain at 120V for 60 minutes.

In some cases, PCR products were sent for Sanger sequencing. PCR reactions were purified using the Monarch® PCR & DNA Cleanup Kit (T1030S, New England Biolabs). Sanger sequencing samples were prepared by combining 150 ng purified PCR product and 0.25 μM forward (or reverse) primer to a final volume of 10 μL. All sequencing was run by staff at the Molecular Biology Service Unit (MBSU) at the University of Alberta. Sequencing results were analyzed with SnapGene View 6.0.2.

### 2.5 Apilimod Treatment

The PIKfyve inhibitor apilimod (SML2974, Sigma-Aldrich) was resuspended in dimethyl sulfoxide (DMSO) to a concentration of 5 mM by gently heating. 1 mM aliquots were stored at −20°C and stock inhibitor was stored at −80°C. Drug exposure occurred in 24-well plates and zebrafish larvae were exposed from 4-6 dpf or 5-7 dpf to concentrations of 100 nM to 1 μM apilimod in EM, with controls exposed to 1% DMSO/EM or just EM. The drug was removed through three EM washes at the termination of the experiment and fish were euthanized as described above.

### 2.6 Electroretinography

Full-field electroretinography (ERG) was used as a non-invasive technique to measure the electrical response of the fish retina to light stimuli, following our previously published protocol (Nadolski et al., 2020). Reference and recording electrodes were prepared by removing 2 cm of insulating housing from a platinum electrical lead wire and intertwining the exposed platinum with 4 cm of 32-gauge silver wire. Both electrodes were placed into household bleach (5.25%) for 5 minutes to increase electrode conductivity and then air dried for three minutes. The reference electrode was positioned on top of a 35 mm petri dish, fitted with a sponge soaked in anesthetic, and secured to a 3D-printed testing platform. Zebrafish larvae were anaesthetized in TMS until unresponsive and transferred to filter paper, which was then placed on the soaked sponge. The tip of the recording electrode was positioned on the centre of the cornea using a micromanipulator. The platform was placed into the ColorDome Ganzfeld light stimulator for ERG testing using the E3 Electrophysiology System. Anesthetized fish were flashed with 10 cd·s/m^2^ light five times and the average response was calculated from the five individual ERG traces. Waveform quantification was done through the system’s built-in algorithm.

### 2.7 Melanin Quantification by Fluorescence Spectroscopy

To analyze whole embryo melanin formation after apilimod exposure, we adapted a previously published fluorescent spectroscopy protocol (Fernandes et al., 2016). Briefly, 56 hpf embryos were euthanized and groups of 65 embryos were lysed by centrifuging at 10,000 g in 1 mL of 10% DMSO/1N NaOH. After centrifugation, the supernatant was discarded and the pellet was resuspended in 500 uL of 10%DMSO/1N NaOH. At this step, the pellet was frozen at −20℃ for later use. Melanin was solubilized by incubating the pellet at 80℃ for 1 hr with occasional vortexing. The solution was centrifuged at 3000 g for 5 minutes and 120 uL supernatant (in replicates of four) was transferred to a PCR tube. 36 uL of 30% H₂O₂ was added to the PCR tube and incubated for 4 hrs at room temperature. Before measuring fluorescence, the solution was centrifuged at 3000 g for 5 minutes and then 150 uL was transferred to a 96-well COSTAR microplate. Melanin fluorescence was measured in quadruplicate using an excitation wavelength of 470 nm and emission wavelength of 550 nm in a COSTAR microplate reader. Experimental group was embryos exposed to 500 nM apilimod in 1%DMSO/EM at 22 hpf. Positive control groups: siblings raised in EM, siblings raised in 1%DMSO/EM. Negative control groups: siblings raised in EM containing 60 mg/L of 1-phenyl-2-thiourea (PTU) to block pigment formation.

### 2.8 Cryopreservation and Cryosectioning

Zebrafish larvae were euthanized in 4.2% TMS and then fixed in 4% paraformaldehyde (PFA) at 4°C for 1-3 days. After fixing, the samples were washed three times with 1X PBS (5 min each) and placed in 17.5% sucrose at 4°C for cryoprotection. Once the eyes sank to the bottom of the tube, the solution was replaced with 35% sucrose and stored at 4°C overnight. Eyes were embedded in cryomolds filled with Fisher Healthcare Tissue Plus O.C.T Compound Clear and stored at −80°C until sectioning. Eyes were sectioned at 12 μm using a Leica cryostat and Fisherbrand Superfrost Plus slides and stored at −20°C until staining.

### 2.9 Paraffin Embedding and Sectioning

Zebrafish 6 dpf larvae were euthanized in 4.2% TMS and fixed in neutral buffered formalin (NBF) for 1-3 days at 4°C on a shaker. Samples were washed 3x in PBS and then embedded in agarose (3 fish/block). The blocks were allowed to solidify at room temperature before being placed back into NBF overnight. The next day, samples were dehydrated through a graded ethanol series (50%, 70%, 90%, and 100%) performed in a Leica Tissue Processor 1020.

Samples were then placed in a 1:1 toluene:ethanol (100%) solution before being submerged in toluene. Samples were then placed in liquid paraffin wax overnight to allow for tissue penetration. The next day, agarose blocks were mounted in paraffin blocks and allowed to solidify at room temperature. A Leica microtome was used to cut 5 μM coronal sections of the central zebrafish eye that were collected on Fisherbrand Plain Premium microscope slides. Slides were incubated at 37°C until staining.

### 2.10 Transmission Electron Microscopy

Zebrafish 6 dpf larvae were euthanized in 4.2% TMS for 15 minutes. Euthanized larvae were fixed in 2.5% glutaraldehyde, 2% paraformaldehyde in 0.1M phosphate buffer, pH 7.2-7.4 for 1-3 days at 4°C on a shaker. Tissue processing was carried out in a fume hood and occurred over three days. On day one, fixative was removed with three 0.1M phosphate buffer washes (10 min each) and samples were stained with 1% osmium tetraoxide in 0.1M phosphate buffer for 1 hour. Samples were washed three times in 0.1M phosphate buffer (10 min each) and dehydrated through a graded ethanol series (15 min each, 50%, 70%, 90%, 100% x 3). A 1:1 mixture of 100% ethanol and Low Viscosity Embedding Media Spurr Resin was used to infiltrate the dehydrated samples for 1-3 hours. Next, samples were left in 100% Spurr resin overnight. On day two, the Spurr resin was changed twice in the morning. Samples were embedded in flat molds with Spurr resin and sample ID paper labels before being cured in a 70°C incubator overnight. On day three, samples were removed from the oven following resin hardening and stored at room temperature until sectioning.

Samples were sectioned by Dr. Kacie Norton at the Biological Sciences Microscopy Unit at the University of Alberta using a Reichert-Jung Ultracut E Ultramicrotome to generate sections of 70 to 90 nm thickness. Sections were stained with uranyl acetate and lead citrate stain. Ultrathin sections were imaged with a Philips – FEI Morgagni 268 transmission electron microscope operating at 80 kV. Images were captured with a Gatan Orius CCD Camera. Pigment granule analysis was performed in ImageJ.

### 2.11 Immunohistochemistry

Cryosections were stained with a variety of primary and secondary antibodies according to the following protocol. If staining occurred immediately after sectioning, slides were allowed to air-dry for at least 2 hours. If slides were frozen after sectioning, then they were allowed to come to room temperature before staining. A lipid line was drawn around sections with a Cole-Parmer Essentials Hydrophobic Barrier PAP Pen. Slides were placed on a rack in a tinfoil-wrapped 150 mm petri dish lined by a moistened Kim wipe. Slides were washed three times (10 minutes each) with 1X PBTD (0.1% Tween-20, 1% DMSO in 1X PBS). Next, 200 μL of block solution (2% goat serum in 1X PBTD) was added and slides were incubated at room temperature for 30 minutes. The block solution was removed and replaced with 200 μL of block solution containing the primary antibodies. A piece of Parafilm was gently placed on top of each slide and the plates were covered and incubated at 4°C overnight. The next day, slides were washed three times in 1X PBTD (10 minutes each), and 200 μL of block solution containing secondary antibodies and/or stains was added. Slides were incubated for one hour at room temperature before being washed three times with 1X PBTD (10 minutes each) and a coverslip applied with Mowiol mounting media (made and kindly provided to us by Dr. Simmonds, University of Alberta) and stored at 4°C until confocal imaging. The following primary antibodies and concentrations were used: mouse monoclonal anti-rhodopsin 4D2 (Novus, US) at 1:500, mouse monoclonal anti-ZPR2 (ZDB-ATB-081002-44, Zebrafish International Resource Center, Oregon, US) at 1:100, rat monoclonal anti-LAMP1 (1D4B, AB_528127, Departmental Studies Hybridoma Bank, University of Iowa, US) at 1:200, rabbit anti-LC3A/B (4108S, New England Biolabs, CA) at 1:100. The following secondary antibodies and concentrations were used: goat anti-mouse IgG Alexa Fluor 680 (A21057, Invitrogen, US) at 1:500, goat anti-rabbit IgG Alexa Fluor 488 (A11008, Life Technologies, US) at 1:500, donkey anti-rat IgG Alexa Fluor 488 (A21208, Invitrogen USA) at 1:500, and donkey anti-mouse IgG Alexa Fluor 594 nm (R37115, Invitrogen USA, gifted by Simmonds Lab, University of Alberta) at 1:500.

### 2.12 Lysotracker Staining

Lysotracker Red DND99 (L7528, Invitrogen, USA) was used to stain lysosomes in live animals. The stain was added to a final concentration of 10 μM to 24-well plates containing six zebrafish larvae in EM. Zebrafish were incubated with the stain at 28.5°C for 2 hours before being rinsed three times with EM to remove the excess stain. Fish were then euthanized in 4.2% TMS and fixed.

### 2.13 Hematoxylin and Eosin Staining

In a fumehood, slides with paraffin sections were first washed twice in toluene (5 minutes/wash) to deparaffinize. Samples were rehydrated through a graded ethanol series (100%, 90%, 70%, 50%) and then washed in distilled water for 2 minutes. Slides were stained with Hematoxylin Gill III (Leica Biosystems) for 2 minutes and then rinsed with cold tap water for 15 minutes to remove the stain. Slides were placed in 70% ethanol for 2 minutes before being stained with Eosin (Leica Biosystems) for 30 seconds. Slides were washed twice in 100% ethanol to remove the eosin stain (2 minutes each) and kept in toluene during coverslipping with DPX mounting media (Electron Microscopy Sciences, Hatfield, USA). Slides were dried in a 37°C incubator overnight and stored at room temperature until imaging.

### 2.14 TUNEL Staining

Cryosections collected on SuperFrost Plus slides were fixed in freshly prepared 4% paraformaldehyde for 20 minutes at room temperature and then washed three times in 1X PBS (10 min each). Slides were incubated for 2 minutes on ice in a permeabilization solution consisting of 0.1% Triton X-100 in 0.1% sodium citrate. Slides were rinsed twice with PBS and then 50 μL of the TUNEL reaction mixture, consisting of 5 μL TUNEL Enzyme (11767305001, MilliporeSigma) and 45 μL TUNEL Label (11767291910, MilliporeSigma), was added to the sections. Slides were incubated for 60 minutes at 37°C in a humid, dark environment before being rinsed three times in PBS and coverslipped with Mowiol mounting media. Slides were stored at 4°C until imaging.

### 2.15 Light and Confocal Microscopy

A Zeiss Axioscope.A1 and a SeBaCam camera (Laxco Inc., Mill Creek, WA) was used to image histological retinal sections. A Zeiss LSM 510 Meta Confocal Microscope was used to image fluorescent tissue sections. Images were processed using ImageJ. The Analyze Particles tool was used to assess melanosomes.

### 2.16 Statistical Analysis

All statistical analysis was performed in GraphPad Prism Version 9.3.0. Where comparisons were between two groups, an unpaired t-test was performed. Where comparisons between more than two groups were performed, a one-way ANOVA followed by Tukey’s posthoc test was performed. An outlier test was done to remove outliers from the data. All measurements are reported as mean and standard deviation.

## 3. RESULTS

### 3.1 CRISPR mutagenesis-mediated loss of PIKfyve leads to early lethality

The zebrafish genome contains a single orthologue of the human *PIKFYVE* gene, known as *pikfyve* and located on chromosome 9. The gene is highly conserved between zebrafish and humans, sharing 71% sequence identity between the longest transcripts. *pikfyve* is broadly expressed in the developing and adult zebrafish neural retina, with RPE expression unreported due to pigmentation (Boisset et al., 2008). We designed CRISPR RNAs (crRNAs) to target two exons encoding a portion of the PIP kinase domain and in the approximately equivalent location of *PIKFYVE* mutations identified in a family with congenital cataracts (Mei et al., 2022). The crRNAs were combined with tracrRNA and Cas9 Alt-R protein and injected into single-cell zebrafish embryos (Hoshijima et al., 2019). Over 30% of embryos injected with the *pikfyve* CRISPR died between 5 and 7 dpf (Fig. S1A). Larval death likely resulted from loss-of-function *pikfyve* mutations given that the timeline matches the early lethality previously described for *pikfyve* knockouts (Mei et al., 2021). As a control for the CRISPR injection procedure, zebrafish were injected with two crRNAs targeting the pigment gene *slc45a2*, demonstrating successful gene disruption and normal survival compared to uninjected siblings.

From the 5-10% of *pikfyve* CRISPR-injected fish that survived to adulthood, we were unable to identify carriers to create a stable heterozygous line. Instead, we analyzed the injected fish (referred to hereafter as *pikfyve* crispants), which would be expected to carry a variety of mutations in the target region. The high death rate between 5 and 7 dpf suggested highly efficient mutagenesis, and indeed, we discovered consistent phenotypes during the subsequent analysis.

To further validate CRISPR-mediated mutagenesis in the crispants, we PCR amplified a genomic region spanning the targeted exons. When the PCR products were run on an agarose gel, two bands were apparent in three of four samples, with the smaller band consistent with the expected amplicon size if the sequence between the two targeted cut sites was removed (∼370 bp, Fig. S1B). Sanger sequencing of the bands revealed the presence of variable sequence starting close to the first expected cut site (Fig. S1C).

Along with increased mortality, *pikfyve* crispants exhibited a mild but significant decrease in body length and variable levels of edema (Fig. 2). Further, *pikfyve* crispant fish failed to develop a swim bladder, which may have contributed to the high larval mortality rate.

**Figure 2.**
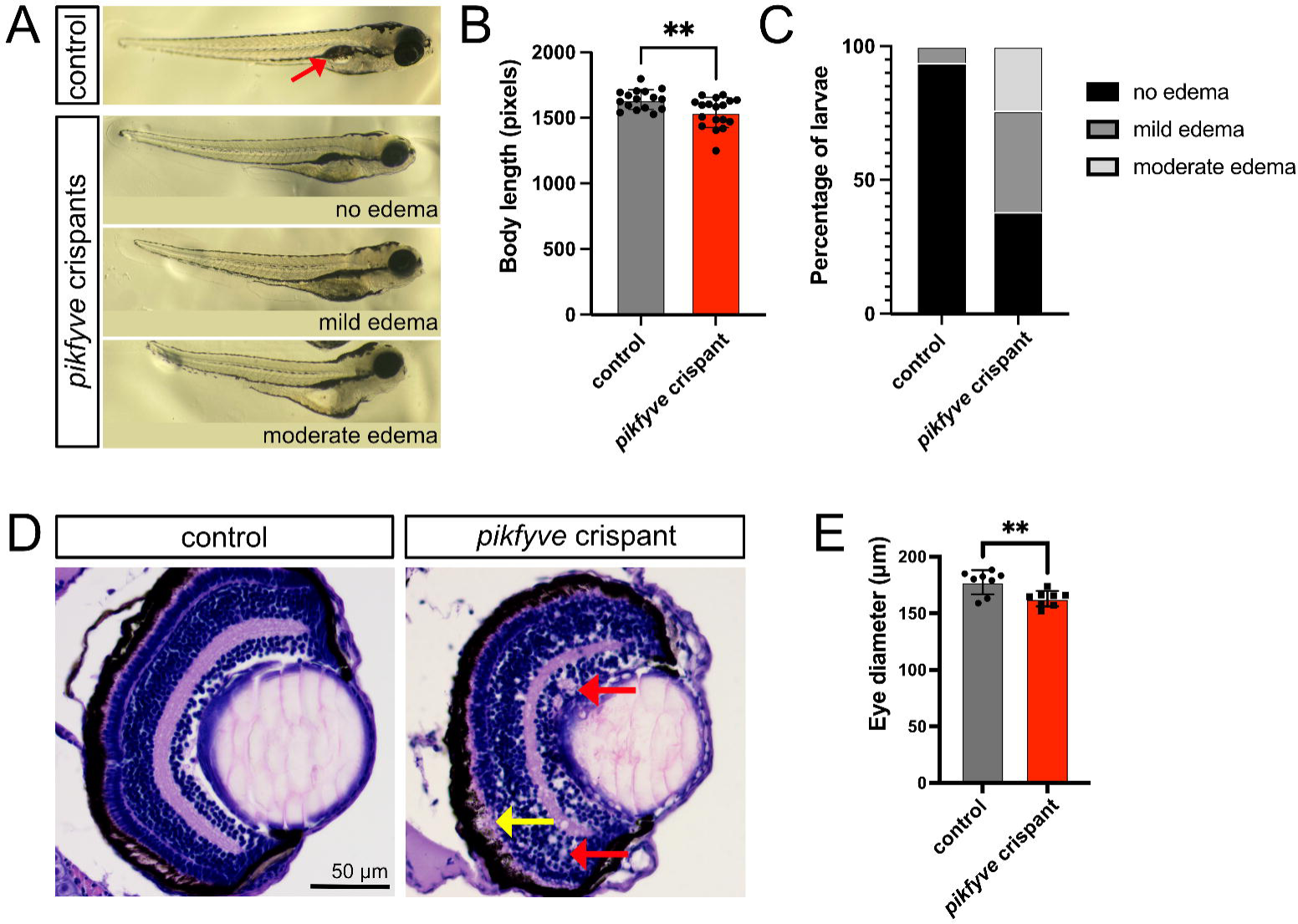
*pikfyve* crispant zebrafish larvae exhibit gross morphological changes. A) Brightfield stereomicroscope images of 6 dpf uninjected control and *pikfyve* crispant embryos, demonstrating that many of the crispant larvae exhibit a shorter body length, edema, and failure to inflate swim bladder (red arrow shows inflated swim bladder in control fish). B-C) Quantification of body length and edema in *pikfyve* crispants (n=18 larvae) and uninjected controls (n=16 larvae). D) H&E stained paraffin sections of eyes from uninjected control and *pikfyve* crispant 6 dpf zebrafish larvae. Red arrows point to vacuoles in crispant retina and yellow arrow to gap between RPE and outer nuclear layer (ONL). E) Quantification of eye size for uninjected control (n=9 larvae) and *pikfyve* crispant fish (n=8 larvae). **p<0.01

### 3.2 Retinal organization is generally maintained in *pikfyve* crispants

Eye architecture was analyzed in H&E-stained paraffin sections of 5 dpf *pikfyve* crispants and control fish. The *pikfyve* crispant eyes were significantly smaller than uninjected siblings, consistent with the reduced body size (Fig. 2D, E), and vacuoles were present around the lens as reported previously in *pikfyve* mutants (Mei et al., 2022). General retinal organization was maintained in the crispants, with the expected three neural layers separated by two plexiform layers, as well as an intact RPE. However, the layers were more variable in thickness and appeared less orderly. In three of the eight *pikfyve* crispant eyes analyzed, there was a region with abnormal expansion between the RPE and ONL, in the area of the photoreceptor OSs (Fig. 2D). Further, we observed vacuoles present across multiple retinal layers in five of the eight eyes.

### 3.3 Accumulation of vacuoles in RPE and photoreceptors of *pikfyve* crispants

Next, we utilized TEM to visualize retinal ultrastructure, specifically focusing on the RPE and photoreceptors of 6 dpf *pikfyve* crispants and uninjected control zebrafish (Fig. 3). We observed profound differences between *pikfyve* crispants and controls. There was a striking accumulation of vacuoles in the crispant RPE, resulting in an inconsistent thickness of the RPE layer and areas of massive expansion (Fig. 3A). Many of the vacuoles appeared fluid-filled and lacked obvious internal structure, while others contained melanosome pigment. Notably, the RPE of both control and crispant fish contained only a few isolated phagosomes, and therefore we did not detect evidence of stalled phagocytosis using the crispant model of PIKfyve loss.

**Figure 3.**
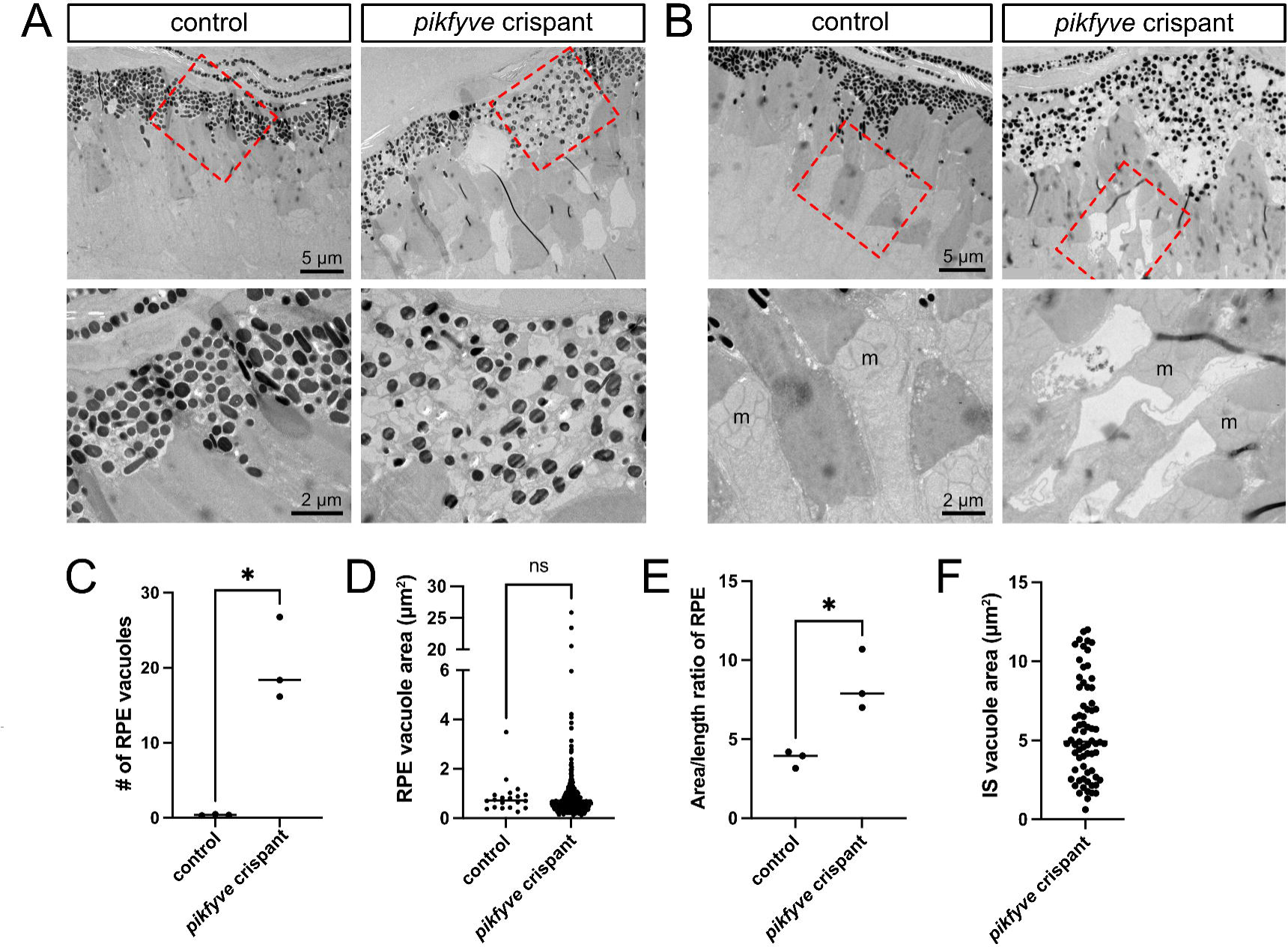
Ultrastructure of outer retina in *pikfyve* crispants reveals vacuole and phagosome accumulation. A-B) TEM images of 6 dpf uninjected control and *pikfyve* crispant zebrafish larvae, highlighting the buildup of vacuoles in the RPE of crispants, accompanied by expansion of the RPE (A), and the presence of large vacuoles in the crispant photoreceptor inner segments, just below the mitochondrial (m) cluster (B). C) Quantification of average number of vacuoles per 10 µm length of RPE in control uninjected and *pikfyve* crispant larvae. Each point represents an individual fish, averaged across multiple sections (n=3 fish per group). D) Average area of RPE vacuoles for 3 fish per group. E) Quantification of area/length ratio of RPE provides measure of RPE expansion in *pikfyve* crispants compared to control fish. Each point represents the ratio for an individual fish, averaged across multiple sections (n=3 fish per group). F) Inner segment (IS) vacuole area, pooled values measured in sections of retinas from three *pikfyve* crispant larvae.

Photoreceptors also consistently developed vacuoles in the *pikfyve* crispants, although with different characteristics than observed in the RPE. Crispant photoreceptors typically contained a single large vacuole located in the inner segment between the nucleus and mitochondrial cluster, and containing fragmented membranous material (Fig. 3B).

RPE vacuole number was quantified in Fig. 3C, showing a significant increase in the *pikfyve* crispants. RPE vacuole area was also quantified and although an occasional large vacuole was observed in the crispants, the vast majority of the vacuoles were less than 1 µm^2^ and therefore the average vacuole size was comparable to the occasional vacuoles observed in control RPE (Fig. 3D). The average RPE thickness, measured as the ratio of the RPE area to length, was more than doubled in the crispants (Fig. 3E). Inner segment vacuoles in the crispants had a mean area of 5.6 ± 3.1 μm^2^, while no vacuoles were detected in photoreceptors of control fish (Fig. 3F).

### 3.4 Lack of visual responses in pikfyve crispants

In order to evaluate vision in the *pikfyve* crispants, we measured the electrical response of the retina to flashes of light using our previously-published zebrafish electroretinography (ERG) procedure (Nadolski et al., 2020). All fish were evaluated in light-adapted conditions and recorded responses were therefore primarily cone-mediated. Eleven *pikfyve* crispant fish underwent ERG testing and nine of the fish exhibited flat ERGs with no significant a-wave or b-waves recorded (Fig. 4A-C). The a-waves and b-waves for *pikfyve* crispants were quantified by the program’s built-in algorithm, which picks the lowest and highest peaks within a predetermined timeframe and assigns them as the a-wave and b-wave, respectively. Nevertheless, it is difficult to analyze a-wave and b-wave implicit time for the *pikfyve* crispants as there was no significant wave recorded. An outlier test was performed on the b-wave data and the two *pikfyve* crispant fish that showed measurable responses were removed. These two fish exhibited wildtype-like responses, suggesting unsuccessful CRISPR mutagenesis of *pikfyve*. All eight of the uninjected siblings showed significant responses to light with little variation in implicit time of a-waves and b-waves and some variation in the amplitudes of the two waves. These findings suggest that *pikfyve* plays a critical role in development of visual function.

**Figure 4.**
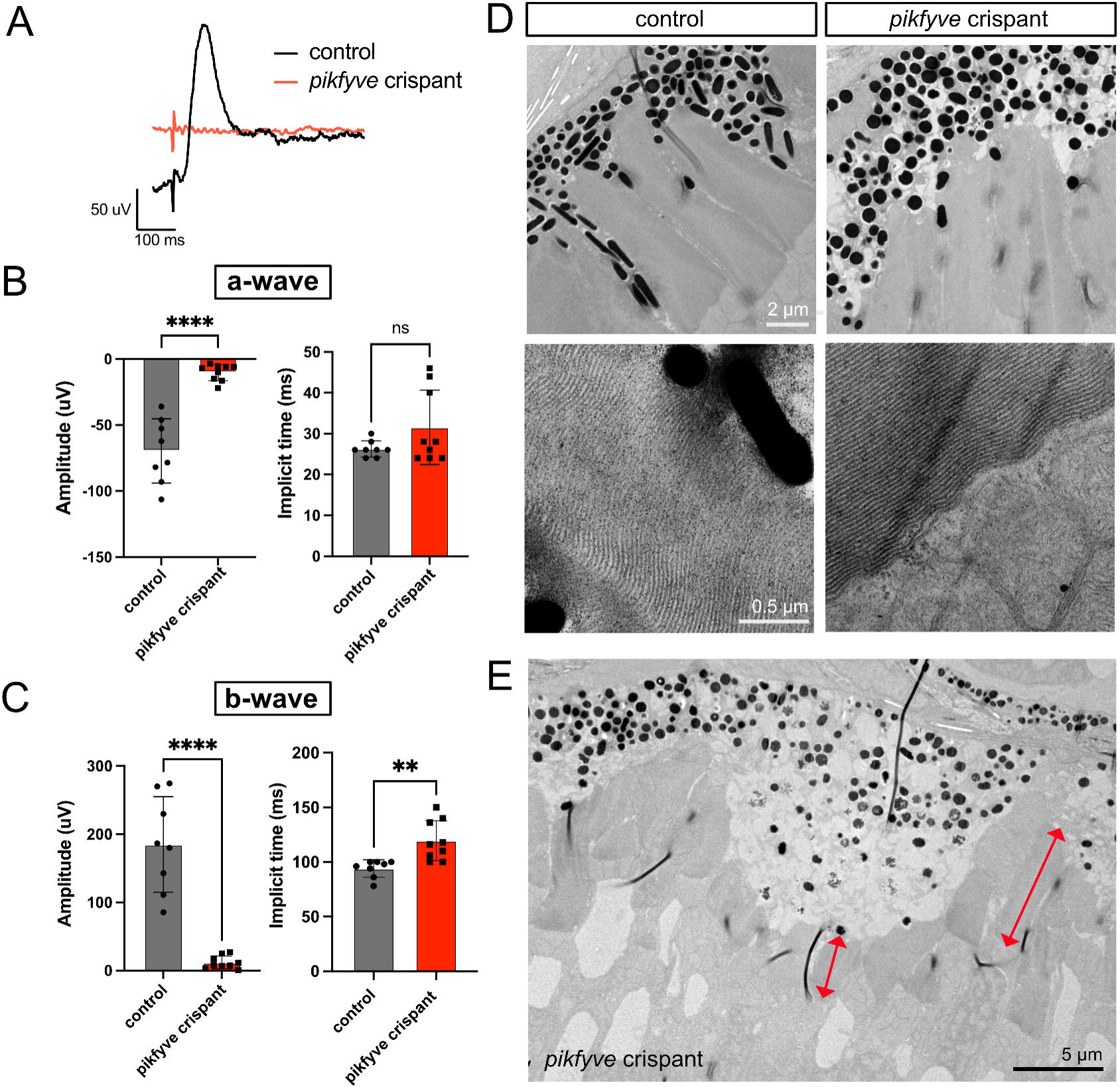
Absent visual responses of *pikfyve* crispants despite largely preserved outer segments. A) Representative photopic electroretinography (ERG) traces from *pikfyve* crispant and uninjected control zebrafish at 5 dpf. B) Quantification of a-wave amplitude and implicit time. Note that the implicit time was difficult to assess in the *pikfyve* crispants given the lack of a visible a-wave. C) Quantification of b-wave amplitude and implicit time. Each dot on the graphs in B-C represents the average measurement of five recordings from an individual fish (n=8 uninjected controls, n=9 *pikfyve* crispants). F) TEM images of photoreceptor outer segments (OS) in 6 dpf control and *pikfyve* crispant fish reveals no apparent disorganization of OS discs in crispants. G) In areas of RPE expansion in *pikfyve* crispants, OS length is reduced (red double arrows indicate OS length for two photoreceptors).

Given the profound loss of vision, we examined the photoreceptor OS in the crispants. The OS discs appeared properly formed with layering indistinguishable from controls (Fig. 4D). However, photoreceptors underlying regions of significant RPE expansion exhibited shortened OS (Fig. 4E). Nevertheless, the observed OS changes are unlikely to explain the full vision loss in the *pikfyve* crispants.

### 3.5 PIKfyve disruption after embryonic development impairs RPE and photoreceptor homeostasis

In order to differentiate between the role for PIKfyve in retinal development versus ongoing cellular function, we created an assay for blocking PIKfyve in larval fish using the specific inhibitor apilimod. Treatments were done over two days, starting at 4 or 5 dpf and thereby allowing normal embryogenesis to occur prior to inhibition. Exposure to 1 uM or above was lethal and therefore doses of 100 to 500 nM were used.

PIKfyve function in differentiated photoreceptors and RPE was first assessed by analyzing TEM images of zebrafish retinas following treatment with apilimod or vehicle control from 4-6 dpf (Fig. 5). Similar to *pikfyve* crispants, apilimod-treated fish exhibited an RPE layer of inconsistent thickness, with massively expanded regions extending into the photoreceptor layer (Fig. 5A). Average RPE thickness was approximately doubled with loss of PIKfyve function (Fig. 5C). Vacuole accumulation was notable, but we also observed a significant number of phagosomes containing OS discs, variably localized from apical to basal (Fig. 5B, D-E). The RPE of control fish rarely displayed disc-containing phagosomes, as expected given that the fish were euthanized approximately 4 hours post-light onset and OS tip engulfment in zebrafish occurs primarily in bursts following the transition to light or dark, with phagosome degradation occurring within an hour (Moran et al., 2022). The presence of disc-containing vacuoles in the *pikfyve* crispant RPE suggests the engulfment of OS discs is uninterrupted, but their degradation is impaired.

**Figure 5.**
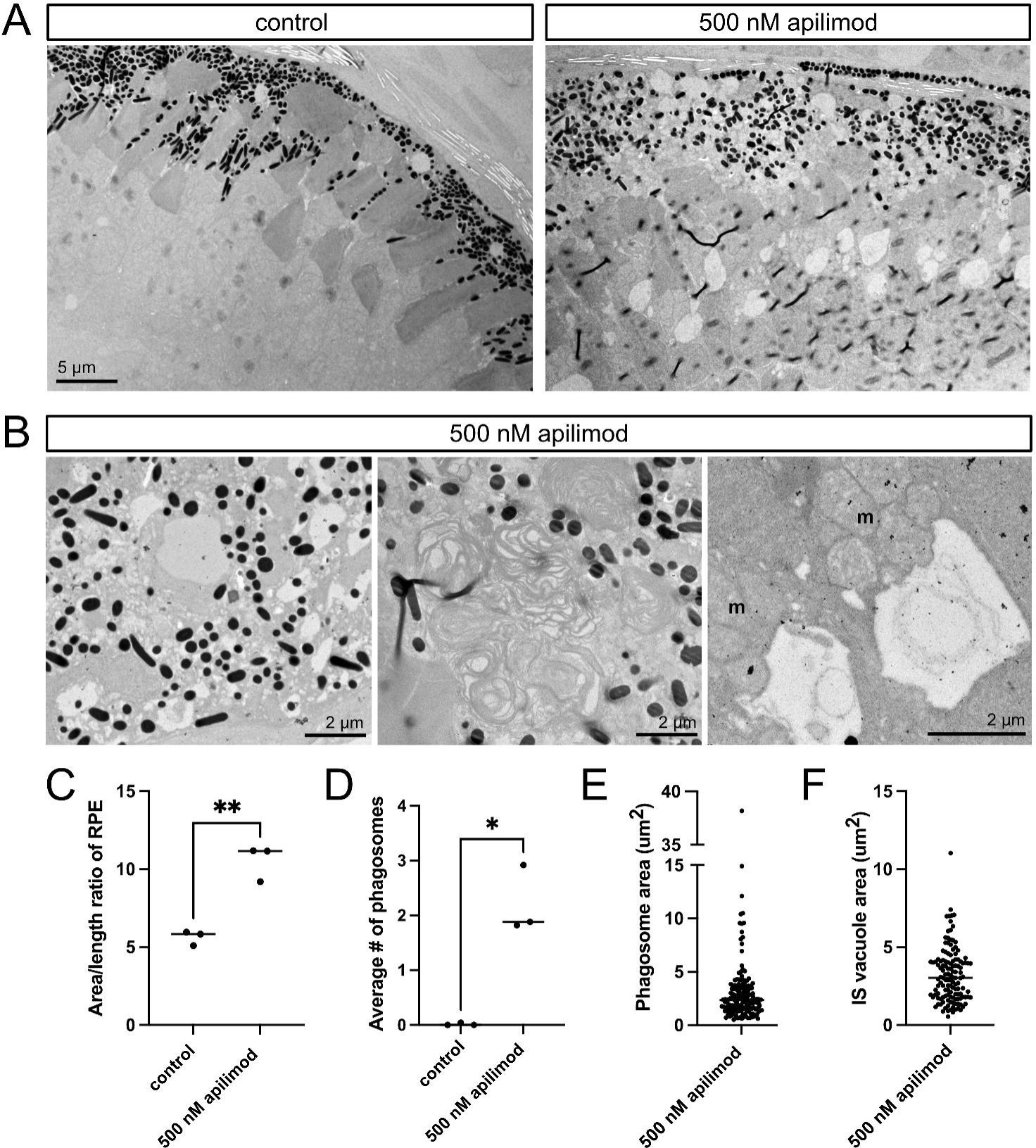
PIKfyve inhibition post embryogenesis induces outer retina vacuolation and impaired phagocytosis. A) TEM images of outer retina from zebrafish larvae exposed to 1% DMSO in embryo media (controls) or 500 nM apilimod from 4 to 6 dpf. B) Magnified TEM images of zebrafish larvae exposed to 500 nM apilimod showing examples of RPE vacuolation (left), a stalled and expanded RPE phagosome (middle), and large vacuoles in the photoreceptor inner segments, just below the mitochondrial cluster (m). C-F) Quantification based on TEM imaging of zebrafish larvae exposed to 1% DMSO in embryo media (control) or 500 nM apilimod from 4 to 6 dpf. C) Quantification of RPE expansion, measured as area/length ratio. Each dot represents the average measurement for an individual fish (n=3 fish per group). D) Quantification of phagosome number per 10 µm length of RPE. Each dot represents the average measurement for an individual fish (n=3 fish per group). E) Plot of phagosome area, with each dot representing an individual phagosome. Measurements are pooled from three apilimod-treated fish. F) Plot of inner segment vacuole area, with each dot representing an individual vacuole. Measurements are pooled from three apilimod-treated fish. *p<0.05; **p<0.01

Photoreceptor homeostasis was also disrupted by PIKfyve inhibition. The photoreceptor inner segments of apilimod-treated fish frequently had a large vacuole located between the nucleus and mitochondrial cluster and containing membranous inclusions, a phenotype very similar to that observed in the *pikfyve* crispants (Fig. 5B, right panel, Fig. 5F).

To evaluate the effect of post-embryonic PIKfyve inhibition on vision, we conducted photopic ERG on 6 dpf zebrafish following 48-hour exposure to 500 nM apilimod or DMSO control media (Fig. 6). While measurable signals were recorded from both experimental and control fish, fish with inhibited PIKfyve showed consistently impaired responses to light. The a-wave of apilimod-treated fish was significantly diminished and delayed compared to the DMSO control fish, indicating delayed and reduced hyperpolarization of cone photoreceptors in response to light. The b-wave of apilimod-treated fish was also delayed and diminished compared to controls, indicating impaired bipolar cell responses.

**Figure 6.**
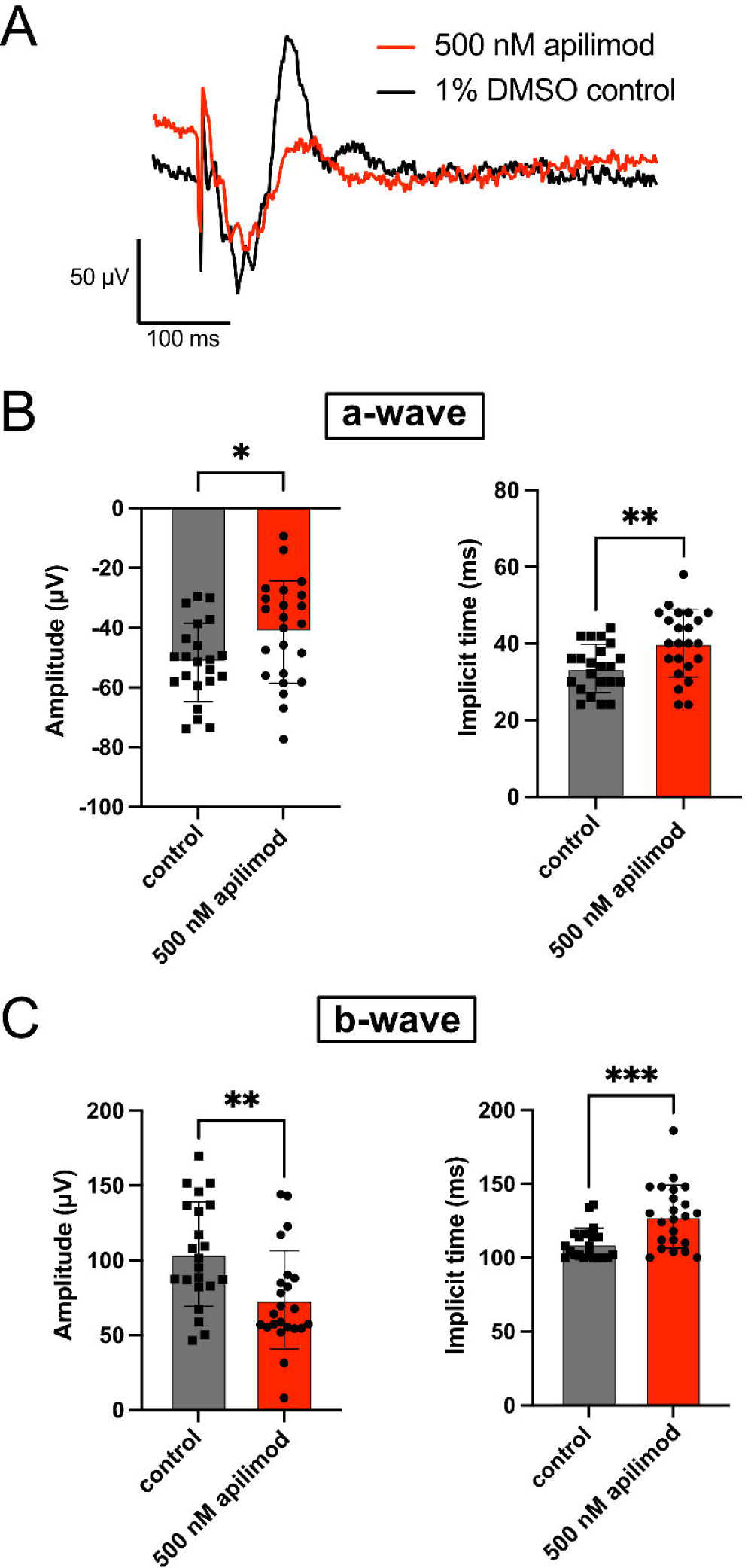
Decrease in visual function following post-embryogenesis inhibition of PIKfyve. ERG recordings were performed on 6 dpf zebrafish exposed to 1% DMSO in embryo media (controls) or 500 nM apilimod from 4-6 dpf. A) Example ERG traces from one control and one apilimod-treated fish. B) Quantification of a-wave amplitude and implicit time. C) Quantification of b-wave amplitude and implicit time. Each dot on the graphs in B-C represents the average measurement of five recordings from an individual fish. n=22 control fish, 23 apilimod-treated fish. *p<0.05; **p<0.01; ***p<0.001

### 3.6 RPE vacuoles in apilimod-treated larvae

To characterize the nature of the vacuoles produced in the RPE upon PIKfyve inhibition, we immunostained retinal sections of *crystal* fish treated with apilimod or control media (Fig. 7). Crystal fish are homozygous for loss-of-function mutations in three pigment genes, and the lack of RPE pigment enables better visualization of vacuoles; however, the fish are more vulnerable to insult and therefore we modified the protocol for a 5 to 7 dpf exposure to improve survival rates. Sections were labelled with antibodies against microtubule light chain 3 (LC3) and lysosomal associated protein 1 (LAMP1), and counterstained with Zpr2 antibody as an RPE marker. LAMP1 marks lysosomes and mature vesicles in the endolysosomal pathway (Cheng et al., 2018; Shearer & Petersen, 2019). LC3 is typically considered a marker of autophagosomes; however, phagosomes can also acquire LC3 in a process termed LC3-associated phagocytosis (LAP), which is active in the RPE (Frost et al., 2015; J.-Y. Kim et al., 2013; Muniz-Feliciano et al., 2017; Peña-Martinez et al., 2022; Tan et al., 2023). Figure 7 shows the presence of LC3 and LAMP1-positive vacuoles in the RPE of apilimod-exposed larvae, the number increasing with higher doses. The vacuoles accumulate distal from the photoreceptor layer, which together with the positive LC3 and LAMP1 staining suggests that these are swollen vesicles in later stages of maturation. Note, photoreceptors are brightly labelled by both antibodies in control and treated conditions, and therefore inner segment vacuoles were not assessed.

**Figure 7.**
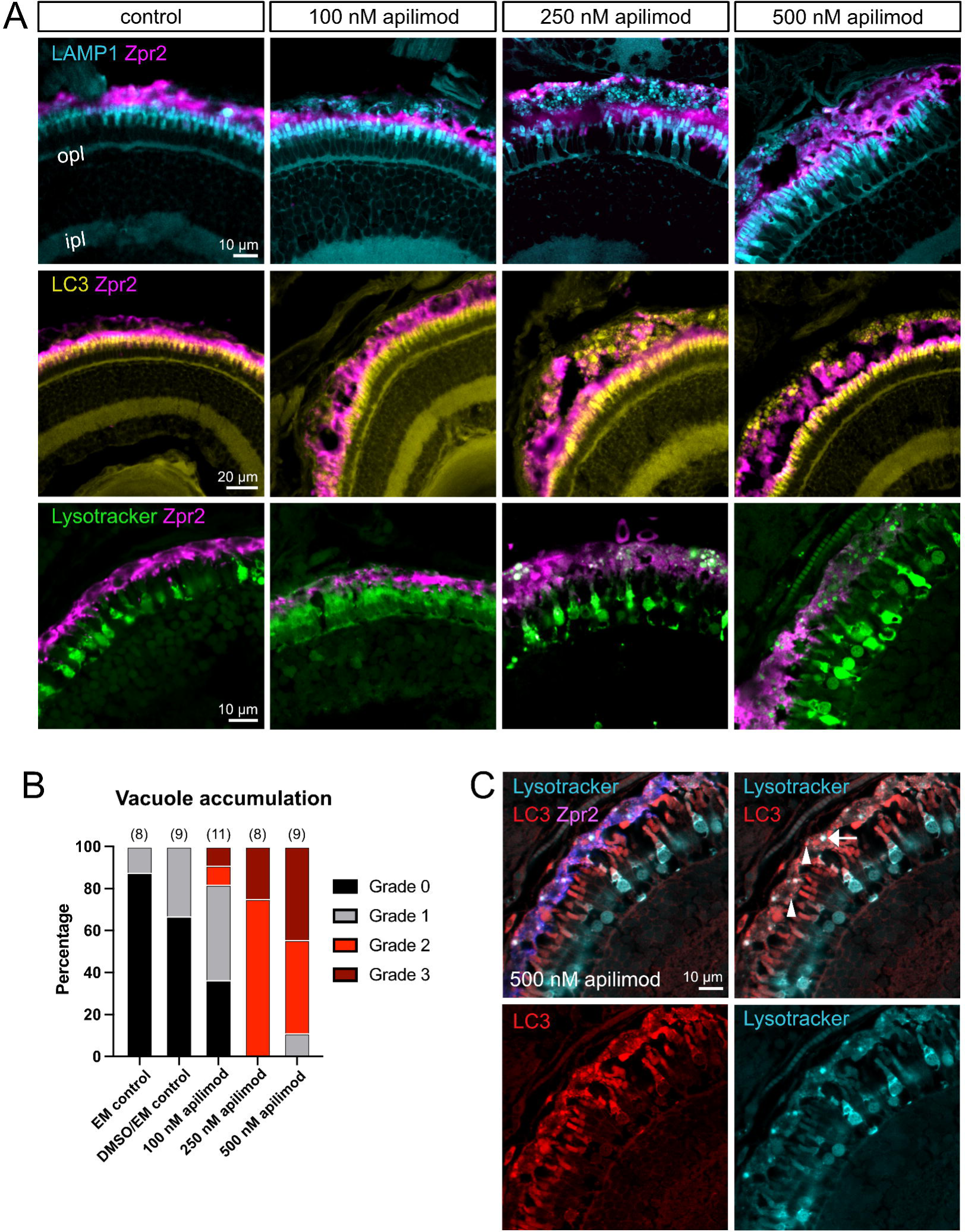
Characterization of enlarged RPE vacuoles after PIKfyve inhibition. Zebrafish *crystal* larvae were exposed to a control solution (embryo media (EM) or 1% DMSO in EM), 100 nM apilimod, 250 nM apilimod, or 500 nM apilimod from 5 to 7 dpf. A) Cryosections were labelled with Zpr2 to highlight the RPE and with anti-LC3A/B antibody or anti-LAMP1 antibody. For a third group, the fish were incubated with Lysotracker Red for 2 hours prior to fixation. B) Quantification of vacuole accumulation in the RPE. Retinal sections were qualitatively ranked according to number and size of vacuoles in the RPE, from no vacuoles (grade 0) to presence of large vacuoles and/or >25 small vacuoles (grade 3). C) Co-staining of RPE vacuoles with LC3 and Lysotracker reveals that only a subset of vacuoles is labelled with Lysotracker. Arrow indicates vacuole co-labelled with Lysotracker and LC3 and arrows indicate LC3+ vacuoles not labelled by Lysotracker. opl, outer plexiform layer

As it was difficult to precisely count or measure individual vacuoles, an observer unaware of the experimental conditions categorized the immunostained retinas by grade according to prevalence and size of vacuoles (Fig. 7B): absence of vacuoles (Grade 0), limited small vacuoles (<10 vacuoles, grade 1), multiple small to medium sized vacuoles (10-25 vacuoles, grade 2), large vacuoles and/or >25 vacuoles (grade 3). While vesicle formation is expected in control RPE as part of physiological endocytosis, phagocytosis, and autophagy, the apilimod-exposed groups had a noticeable accumulation of enlarged vacuoles, consistent with the TEM data. The distribution in phenotypes displayed a dose-response from 100 to 500 nM, although with a notably large difference between the 100 and 250 nM doses: the phenotype of the 100 nM apilimod group was mild, while the 250 nM group was almost indistinguishable from the 500 nM group.

PIKfyve is implicated not only in maturation of vesicles prior to fusion with lysosomes, but also in lysosomal homeostasis, acidification, and resolution. We used Lysotracker to mark acidic compartments and observed that a subgroup of RPE vacuoles were strongly labelled, indicating that acidification was still possible (Fig. 7A). However, many of the vacuoles were not labelled with Lysotracker, which could indicate stalling prior to fusion with lysosomes or insufficient acidification (Fig. 7C).

### 3.7 Cell death and RPE fragility following PIKfyve loss

We observed during retinal sectioning a tendency in the *pikfyve* crispant and apilimod-treated fish for the RPE to split, introducing large holes or almost fully separating the RPE into two layers. Tearing was analyzed in Zpr2-stained sections of apilimod or control-treated larvae and qualitatively assessed by an independent observer (Fig. 8A-B). The degree of tearing increased with the dose of apilimod, thus mirroring the accumulation of vacuoles.

**Figure 8.**
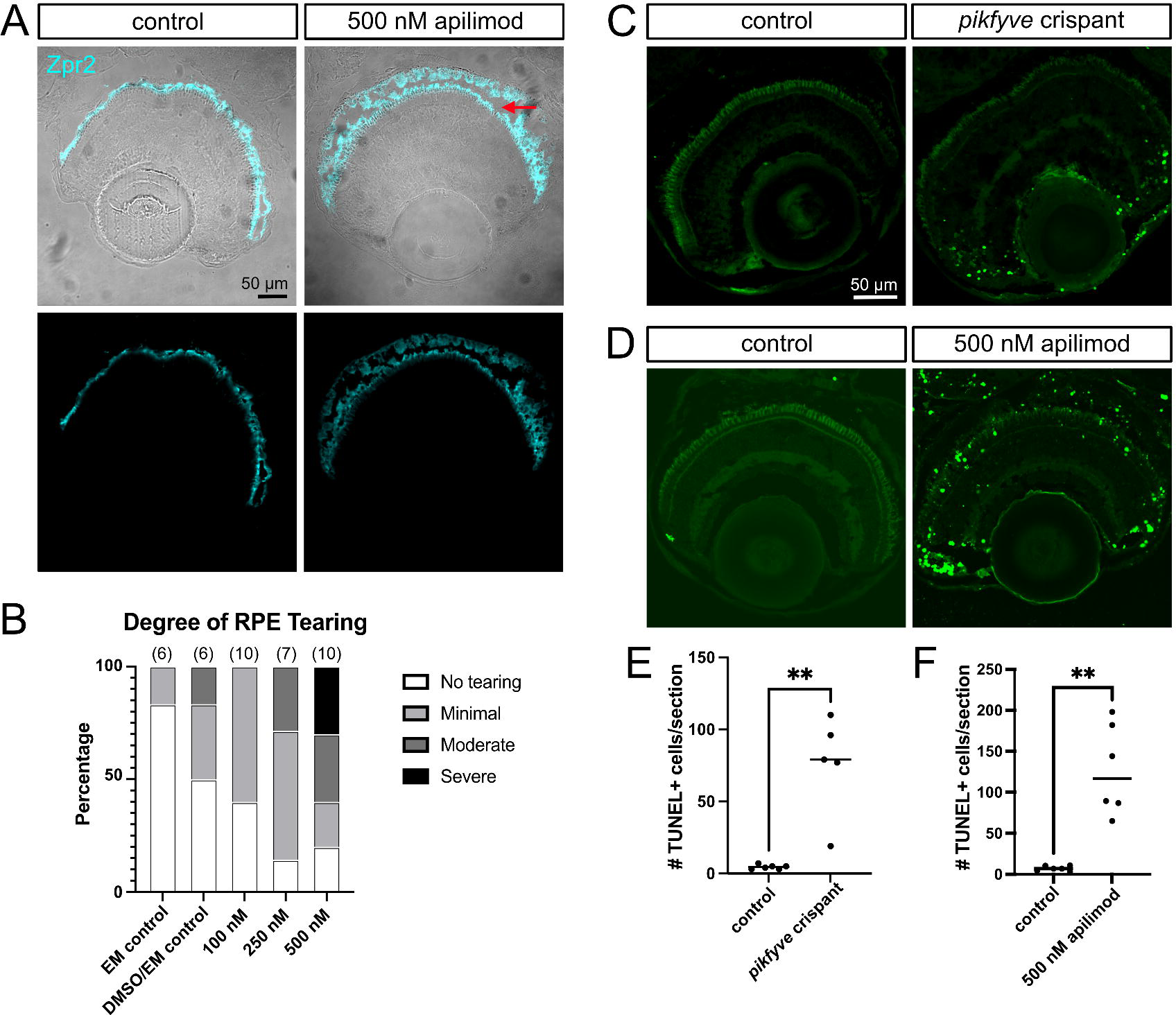
PIKfyve loss leads to fragility of the RPE and cell death in the retinal periphery. A) Examples of retinal cryosections from 7 dpf larval zebrafish exposed to control solution (1% DMSO in EM) or 500 nM apilimod from 5 to 7 dpf. Zpr2 marks the RPE. Arrow highlights tear in RPE of apilimod-treated fish. B) Qualitative analysis of tearing in RPE according to dose of apilimod from 5 to 7 dpf. Control fish were kept in EM or EM containing 1% DMSO. n=# of eyes. C) Images of TUNEL-labelled retinal cryosections from control and *pikfyve* crispant fish at 6 dpf. D) Images of TUNEL-labelled retinal cryosections from fish exposed to 1% DMSO in EM control solution or 500 nM apilimod from 5 to 7 dpf, while exposed to constant light. E) Quantification of average # of TUNEL-positive cells per retinal section for *pikfyve* crispant and control larvae. Each dot represents an individual fish. F) Quantification of average # of TUNEL-positive cells per retinal section for 500 nM apilimod-treated and control larvae. Each dot represents an individual fish. **p<0.01, ****p<0.0001

Although the fragility of the RPE layer could be explained by the accumulation of vacuoles leading to RPE expansion and weakness, cell death may also contribute. We therefore performed TUNEL staining on sections from *pikfyve* crispant and apilimod-treated larvae and appropriate controls (Fig. 8C-D). The apilimod and DMSO-treated fish were additionally stressed by exposure to continuous light for the 48-hour treatment window in order to best capture vulnerable cells. We observed a significant increase in TUNEL-positive cells upon reduced PIKfyve activity, although concentrated in the retinal periphery and overlapping the proliferative tissue of the ciliary marginal zone (CMZ) (Fig. 8C-F). Isolated TUNEL-positive cells were also found in the post-mitotic retina of the apilimod-treated larvae, particularly in the photoreceptor layer. PIKfyve loss can therefore lead to cell death in the retina, with proliferative cells being the most vulnerable followed by photoreceptors. We did not observe apoptosis in the RPE layer and conclude that the tearing is mostly secondary to tissue swelling upon vacuole accumulation.

### 3.8 PIKfyve loss impairs but does not prevent RPE melanosome biogenesis

Melanosomes are lysosome-related organelles that synthesize and store melanin. Melanosome biogenesis occurs in four steps, beginning with stage I melanosomes derived from specialized early endosomes (Bissig et al., 2016; Raposo & Marks, 2007). Previous work implicated PIKfyve at multiple stages of melanosome maturation within skin melanocytes (Bissig et al., 2019; Liggins et al., 2018).

To investigate if PIKfyve is involved in RPE melanosome biogenesis, we inhibited PIKfyve through apilimod treatment of wildtype zebrafish embryos beginning just prior to pigment development (22 hpf) and lasting until 48 dpf, by which time embryos typically exhibit substantial pigmentation. We observed variable and incomplete disruption of both eye and body pigmentation in the apilimod-treated fish at the experiment endpoint, but never a complete lack of pigment comparable to that observed in fish treated with the tyrosinase inhibitor PTU (Fig. 9A). A fluorescent spectroscopy assay was used to quantitatively measure melanin formation in groups of whole embryos. The experimental group was treated with 500 nM apilimod from 22-56 hpf. Positive controls for pigmentation were fish raised in embryo media with or without 1% DMSO and the negative control (absent pigmentation) was a group of fish treated with PTU. Importantly, the RPE contributes substantially to total body pigmentation at this stage given the large size and dense pigmentation of the eyes. The spectroscopy assay revealed a significant, but only partial, reduction in pigment upon *pikfyve* inhibition (Fig. 9B).

**Figure 9.**
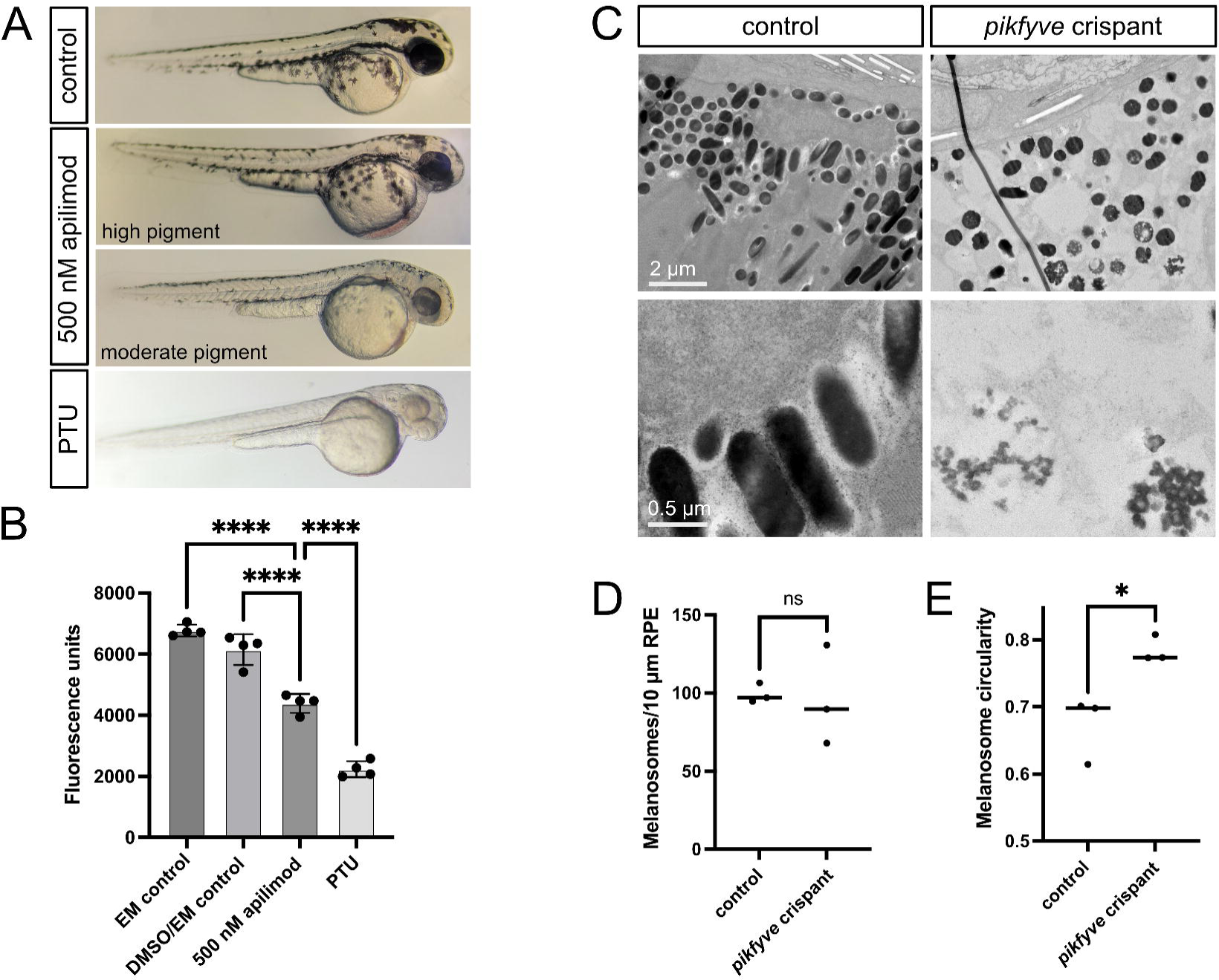
Melanosome biogenesis is mildly disrupted by loss of PIKfyve. A) 48 hpf zebrafish embryos exposed to 500 nM apilimod during melanosome biogenesis (22 to 48 hpf) exhibit a variable reduction, but not total loss, of body and eye pigment. PTU-treated fish is used as a negative control, exhibiting a nearly complete loss of pigment at this stage. B) Melanin quantification by fluorescent spectroscopy showed a significant reduction in pigment in the embryos treated with apilimod from 22-56 hpf compared to fish raised in EM or 1% DMSO in EM, but significantly more pigment than PTU-treated fish. Each dot represents a sample generated from a group of 56 embryos. C) TEM images of 6 dpf control uninjected and *pikfyve* crispant larval zebrafish reveal the presence of fragmented melanosomes and fewer mature rod-shaped melanosomes upon PIKfyve loss. D-E) Quantification of melanosomes in TEM images showing no change in the number of melanosomes per 10 μm length of RPE, but a significant increase in circularity. Dots in D and E represent average measurements from individual fish, n=3 fish per condition. *p<0.05

The mild pigment phenotype suggests that melanosomes can develop and accumulate melanin in the absence of PIKfyve but may not mature correctly. We therefore re-analyzed the TEM images of *pikfyve* crispant fish, where we noted that melanosomes were present in normal numbers within the RPE; however, the pigment was often fragmented and the organelles matured less often into rod-shaped melanosomes. Further, many of the vacuoles in the RPE of both crispant and apilimod-treated fish contained dense pigment regions, thereby indicative of swollen melanosomes (Fig. 9C-E). Our data suggest that PIKfyve is required for RPE melanosome maturation and homeostasis.

## 4. DISCUSSION

PIKfyve is required for the production of PI(3,5)P_2_ and PI5P, low abundance signalling lipids with unknown functions in the retina. We found that loss of PIKfyve consistently leads to the accumulation of vacuoles in photoreceptors and the RPE, although with notably distinct characteristics: RPE cells accumulate numerous vacuoles of varying sizes, while photoreceptors typically form a single large vacuole immediately below the mitochondrial cluster of the inner segment. Despite the defects in the *pikfyve* crispants, their retinas displayed normal layering indicative of proper cell differentiation. Further, much of the phenotype may arise from impaired homeostasis given that the application of apilimod subsequent to cell differentiation leads to massive swelling of the RPE, disrupted phagosome processing, and vacuole formation in photoreceptors.

RPE cells are considered specialized phagocytes, which by definition are phagocytes that are resident to a particular tissue, form tissue-blood barriers, and facilitate transport (Lakkaraju et al., 2020; Penberthy et al., 2018). The RPE endures the highest phagocytic demand in the body, a burden compounded by the accumulation over time of lipofuscin, an autofluorescent and indigestible material primarily formed of vitamin A aldehyde derivatives known as bisretinoids (Sparrow et al., 2012).

RPE cells extend actin-based apical processes that wrap around OS tips to facilitate scission and engulfment, giving rise to nascent phagosomes (Kwon & Freeman, 2020; Lakkaraju et al., 2020; Lieffrig et al., 2023; Umapathy et al., 2023; Zihni, 2025). Following ingestion, phagosomes undergo a complex series of maturation steps that involve changes to membrane content, merging with endosomes, and apical-basal translocation before finally fusing with lysosomes.

Interestingly, a significant proportion of phagosomes in the RPE are processed by the non-canonical pathway of LC3-associated phagocytosis (LAP) (Frost et al., 2015; J.-Y. Kim et al., 2013; Lakkaraju et al., 2020; Muniz-Feliciano et al., 2017; Peña-Martinez et al., 2022; Tan et al., 2023). Microtubule-associated protein 1A/1B-light chain 3 (LC3) is a ubiquitin-like protein well-characterized as an essential autophagy component added to the autophagosome double membrane by lipidation of LC3-I to form LC3-II (Aman et al., 2021; Noda et al., 2009). The hallmark feature of LAP is the addition of LC3 to single-membrane phagosomes (Martinez, 2018). In the RPE, Tan et al. demonstrated that the adaptor protein optineurin recruits LC3, which mediates interactions with microtubules to promote phagosome transport and fusion with lysosomes (Tan et al., 2023). Here, we showed that PIKfyve loss led to the build-up of LC3-positive vacuoles in the RPE, indicative of stalled autophagosomes or LAP phagosomes. The accumulation of phagosomes containing undigested OS disk membranes in TEM images of the apilimod-treated fish supports the concept that at least a subset of LC3-positive vacuoles are LAP phagosomes.

We also noted that the majority of RPE vacuoles in apilimod-exposed larvae were LAMP1-positive. LAMP1 is typically used as a marker for lysosomes, but also accumulates on late-stage endosomes/phagosomes/autophagosomes (Saharan & Kamat, 2023). As only a subset of vacuoles were robustly labelled with Lysotracker, the LC3- and LAMP1-positive vacuoles could be late-stage vesicles prevented from fusing with lysosomes. On the other hand, some studies report impaired lysosome acidification in the absence of PIKfyve, which could imply that a subset of Lysotracker-negative vesicles were enlarged lysosomes or vesicles that had already fused with lysosomes but failed to properly acidify (Banerjee & Kane, 2020).

One significant finding was the lack of stalled phagosomes in the *pikfyve* crispant RPE. We initially hypothesized that the analysis at 6 dpf was too early to capture disrupted phagosome processing. However, RPE phagocytosis can be observed as early as 5 dpf in zebrafish and the apilimod-treated fish analyzed at 6 dpf showed a dramatic accumulation of disc-containing phagosomes (Gómez Sánchez et al., 2023). Therefore, we postulate that the lack of PIKfyve during RPE differentiation in the crispants led to impaired phagosome formation secondary to general disruption of RPE health and function. The stalled phagosomes in apilimod-treated fish are consistent with a role for PIKfyve in phagosome maturation and not during the engulfment stage.

Autophagy is crucial for maintenance of the RPE and photoreceptors, both long-lived cells continuously exposed to high levels of reactive oxygen species (Markitantova & Simirskii, 2025; Santo & Conte, 2021). Loss of key autophagy genes leads to progressive photoreceptor degeneration and increased vulnerability to light-induced damage (Villarejo-Zori et al., 2021). In addition, photoreceptors continually traffic vast quantities of proteins and lipids into the OS and PIKfyve may contribute to the processing of vesicles transported from the Golgi to the OS, as well as to the cellular response to misfolded proteins. As photoreceptors are not phagocytic, the large inner segment vacuoles induced by PIKfyve loss are likely formed via disruption of endosome and autophagosome processing, but why the defect would manifest as a single large vacuole in each cell is unclear. However, the consistent location is logical given that these cellular processes take place in the inner segment and are spatially restricted by the large mitochondrial cluster apically and nucleus basally.

A developmental defect we observed in the RPE was mild disruption of melanosome structure and embryo pigmentation following early PIKfyve reduction. Melanosome biogenesis progresses through four stages beginning with specialized early endosomes and finishing with opaque organelles densely packed with melanin (Bissig et al., 2016; Raposo & Marks, 2007). In the RPE, many of the melanosomes subsequently acquire an elongated rod shape (Burgoyne et al., 2015). Previous research suggested that PIKfyve is required for membrane dynamics and interactions of stage 1 and 2 melanocytes, and thereby necessary for proper assembly of the PMEL amyloid matrix and delivery of tyrosinase for pigment generation (Bissig et al., 2016; Liggins et al., 2018). Consistent with our findings, melanocyte-specific knockout of *Pikfyve* in mice causes a greying of coat color, but not a full loss of pigment (Liggins et al., 2018). We also noticed that many of the RPE vacuoles in apilimod-treated and crispant larvae appeared to be pigment-containing swollen melanosomes, suggesting an additional defect in homeostasis of mature melanosomes.

Although PIKfyve function had not been previously investigated in the retina, the PIKfyve substrate PI3P has demonstrated roles in both RPE and photoreceptors (Wensel, 2020). Depletion of PI3P through a mutation in the PI-3 kinase Vps34 *in vitro* and *in vivo* leads to impaired recruitment of LC3 to phagosomes in mouse RPE, stalling of phagosome processing and the accumulation of membrane aggregates expressing autophagy markers, eventually resulting in RPE death (He et al., 2025). Similarly, conditional loss of Vps34 in mouse rod photoreceptors leads to impaired fusion of endosomes and autophagosomes with lysosomes, accumulation of large vacuoles containing membrane aggregates and progressive loss of all photoreceptors (He et al., 2016), while cone-specific loss of Vps34 causes degeneration of cones without impacting rods (R. V. S. Rajala, 2021). Our work highlights that the conversion of PI3P to PI(3,5)P_2_ is a likely next step in a critical pathway for photoreceptor and RPE function and survival.

Using ERG, we demonstrated that *pikfyve* crispants and apilimod-treated fish exhibit impaired visual function. Surprisingly, the magnitude of vision loss is far more severe for the crispants than for the apilimod-treated fish despite similar histological changes. The crispant phenotype may be enhanced by defective phototransduction, perhaps secondary to a lack of visual pigment, or by anterior ocular defects reducing light penetration.

A hallmark of PIKfyve inhibition is the accumulation of vacuoles, but the mechanism underlying vacuole formation is not fully understood. PI(3,5)P_2_ is generally implicated during maturation, and not formation, of intracellular vesicles, and also in lysosome resolution following fusion (Bissig et al., 2017; Choy et al., 2018; Hasegawa et al., 2017; G. H. E. Kim et al., 2014; Krishna et al., 2016; Rodgers et al., 2022, 2023). As a signalling lipid, PI(3,5)P2 has multiple partners and its actions may vary between cell types or situations, but a common mechanism is regulation of ion channels, including activation of the cation channels mucolipin1 (TRPML1), two pore channel 1 (TPC1), and TPC2, and inactivation of the chloride channel ClC-7 (Chadwick et al., 2021; Davis et al., 2023; Dayam et al., 2015, 2017; Dong et al., 2010, 2010; Fine et al., 2018; Leray et al., 2022; Li et al., 2019; Wang et al., 2012; Wu et al., 2025; Yuan et al., 2024; Zhang et al., 2012). Impaired ion movement can lead to vacuole formation through blocked vesicle fusion, insufficient acidification, or osmotic swelling. Notably mutations in the TRPML1-encoding gene *MCOLN1* cause Mucolipidosis Type IV, which is associated with a variety of ocular defects including retinal dystrophy and RPE changes (Gibson et al., 2022; Smith et al., 2012) and the *Trpml1-/-* mouse model exhibits thinning of the photoreceptor layer and disrupted OSs (Grishchuk et al., 2016).

Apilimod was, or is currently being, tested in clinical trials as a treatment for Crohn’s disease, psoriasis, neurodegeneration, autoimmune diseases, COVID-19 and more based on its ability to block specific interleukins and promote clearance of neurotoxic protein aggregates through PIKfyve inhibition (Babu et al., 2024; Baranov et al., 2020; Billich, 2007; Krausz et al., 2012; Sands et al., 2010; Wada et al., 2012). Results thus far report a reasonable safety profile; however, the well-characterized rapid vacuolization of cells upon PIKfyve inhibition highlights the potential for apilimod treatment to cause significant side effects. The RPE and photoreceptors reside in a high stress environment and yet need to last a lifetime if vision is to be maintained. Our discovery of profound retinal disruption within two days of apilimod treatment suggests that the proposed clinical use of PIKfyve inhibitors should include regular ophthalmic screening.

Notably, a concurrent study by Rajala et al. (2025) using conditional knockout of murine *Pikfyve* in rods and the RPE produced data that complement our findings and highlight that the phenotypes are largely a result of cell-autonomous effects from PIKfyve loss (R. V. Rajala et al., 2025).

## 5. CONCLUSION

In summary, our data shows that PIKfyve is an essential component of the endolysosomal pathway in photoreceptors and the RPE; loss of PIKfyve function induces vacuole formation and disrupts phagosome processing, melanosome biogenesis, and vision. It would be worthwhile to investigate the long-term consequences of mild PIKfyve inhibition on retinal health, the association of *PIKFYVE* mutations with genetic retinopathies, and whether PIKfyve activity declines with age or in retinal degenerative diseases.

## Acknowledgements

We would like to express our gratitude to Dr. Raju V.S. Rajala from the University of Oklahoma for sharing his data about PIKfyve function in the mouse retina and for critically reading our manuscript. We acknowledge the employees of North Campus Animal Services, University of Alberta, for their excellent fish care. We thank Kacie Norton in the Biological Sciences Advanced Microscopy Facility at the University of Alberta for help with preparation and imaging of TEM and paraffin specimens, Dr. Andrew Simmonds for providing reagents and advice, and Dr. Qiumin Tan for access to the Leica CM1520 cryostat. We greatly appreciate the financial support from the Faculty of Medicine and Dentistry (FoMD) through their subsidization of core facilities. K.A. was supported by an FoMD 75th Anniversary Award and I.A. by an Alberta Innovates Summer Research Studentship.

## CRediT Author Contribution Statement

K.Attia. conceptualization, formal analysis, methodology, investigation, visualization, writing – original draft; I.Anjum. formal analysis, methodology, visualization, investigation, writing – review and editing; S.Lingrell investigation, writing – review and editing; M.Benson. conceptualization, writing – review and editing; I.M.MacDonald. conceptualization, writing – review and editing; J.C.Hocking. conceptualization, supervision, resources, formal analysis, writing – original draft.

## Fund Information

This study was supported by the Edmonton Civic Employees Charitable Assistance Fund through a Department of Surgery Research Award and by an NSERC Discovery Grant (RGPIN-2018-05756).

