## Supplemental Data for "PIKfyve is an essential component of the endolysosomal pathway within photoreceptors and the retinal pigment epithelium"

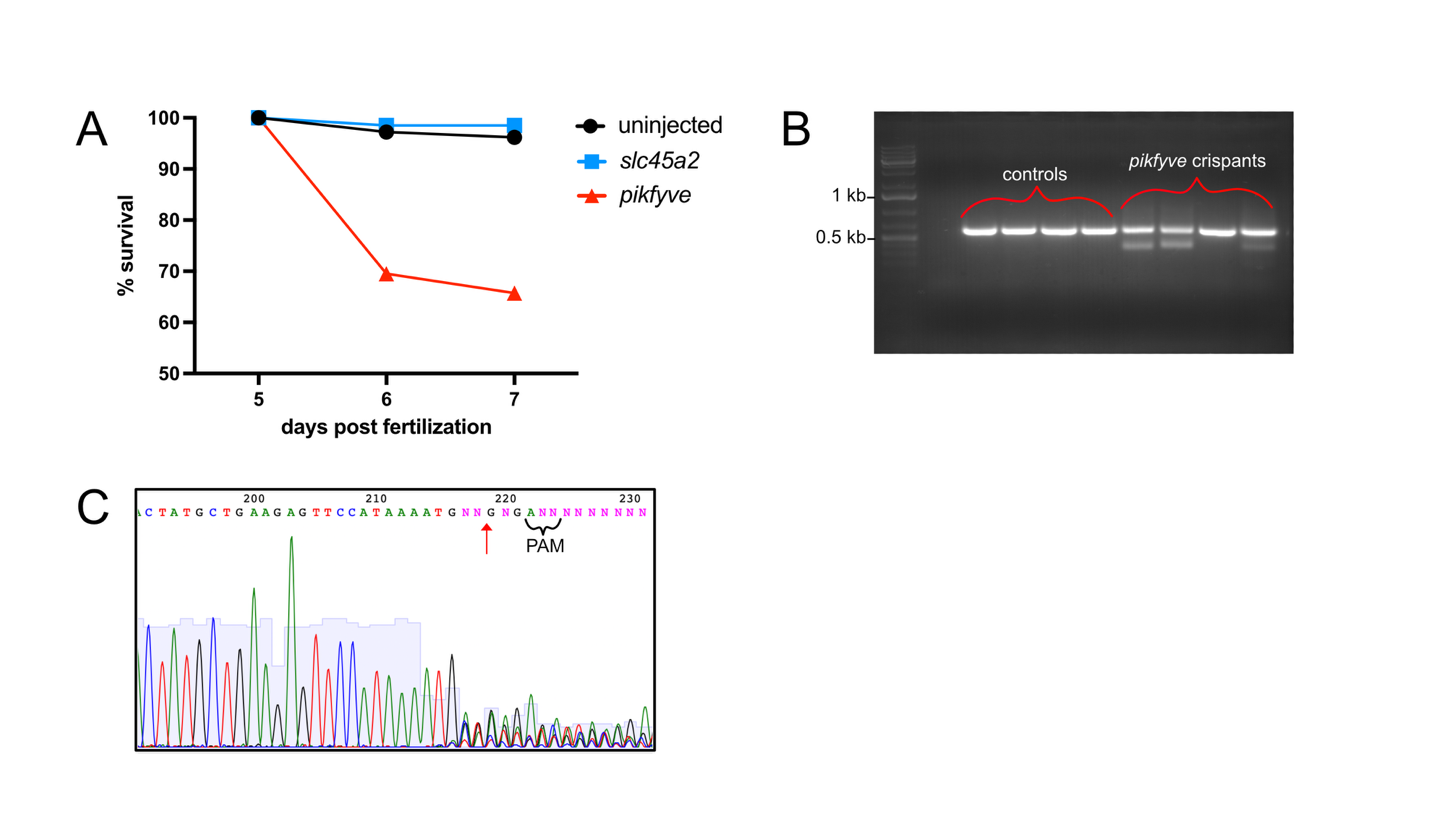

**Figure S1. Injection of CRISPR targeting *pikfyve* in zebrafish causes mutations in the kinase domain and early lethality.** A) Survival curve for uninjected control zebrafish (n=287) and larvae injected with CRISPR targeting *pikfyve* (n=210) or slc45a2 (n=137). B) PCR products from uninjected and *pikfyve* crispant fish. Three of the four *pikfyve* crispant fish have two bands, the smaller being the expected amplicon size (~370 bp) following deletion of the region between the cut sites targeted by the two crRNAs. C) Sanger sequencing of the region targeted by CRISPR, with variable sequence beginning near targeted cut site (red arrow).

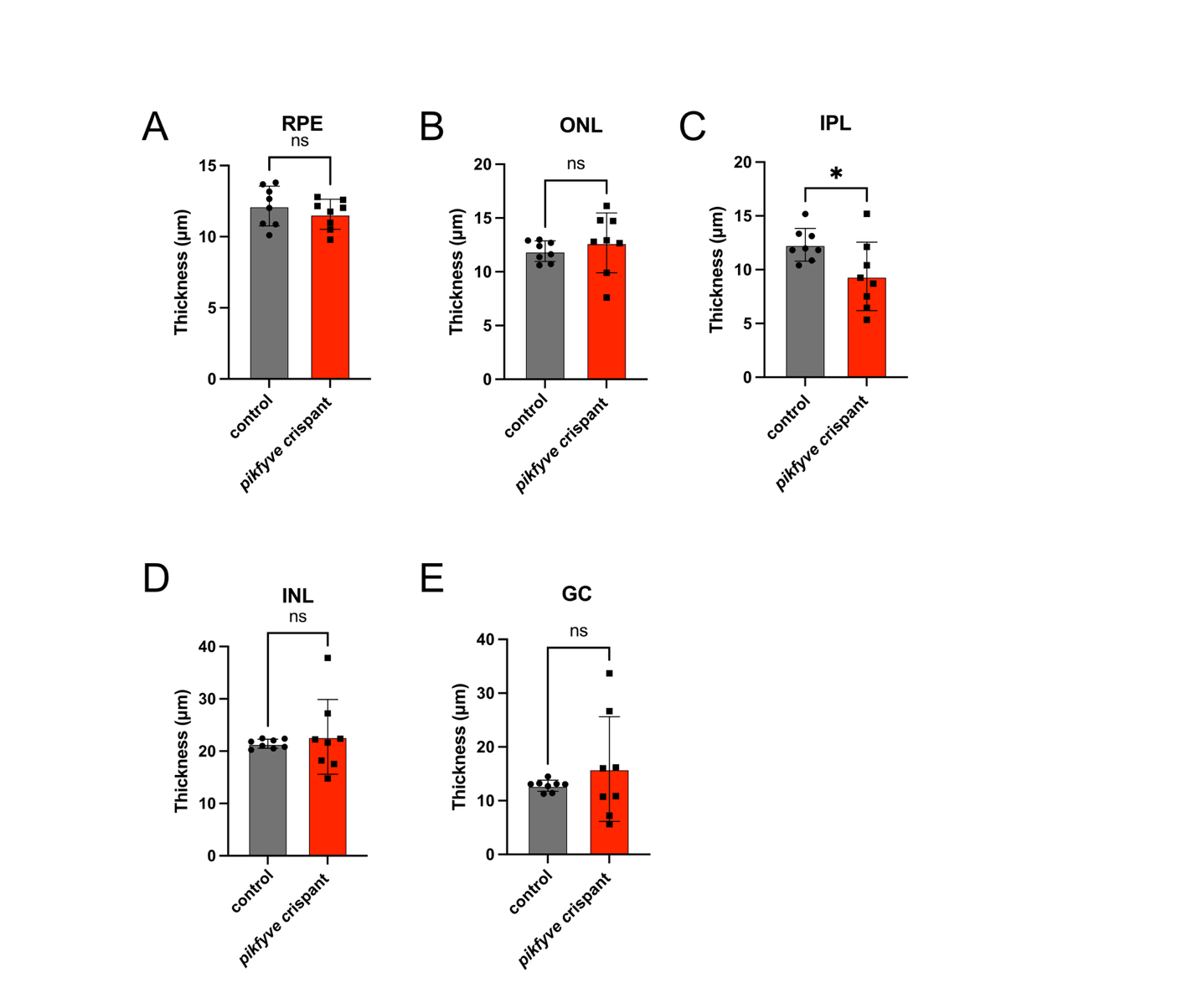

**Figure S2. Quantification of retinal layer thickness in *pikfyve* crispants.** The thickness of retinal layers was measured in H&E sections of 6 dpf *pikfyve* crispants and control uninjected fish**.** RPE, retinal pigment epithelium; IPL, inner plexiform layer; ONL, outer nuclear layer; GCL, ganglion cell layer; INL, inner nuclear layer. *p<0.05; ns, not significant.

| **crRNA** | **Sequence (5’ – 3’)** |
| --- | --- |
| PIK5crExon38 | AGAGUUCCAUAAAAUGCGGGGUUUUAGAGCUAUGCU |
| PIK5crExon39 | UGUGAUGUACGUAAAGUAGUGUUUUAGAGCUAUGCU |
| Slc45a2cr1 | GACGUCUGUACAGUCUGGUGGUUUUAGAGCUAUGCU |
| Slc45a2cr2 | AGUCCAACGCUCAGCAACACGUUUUAGAGCUAUGCU |

**Table S1. crRNA sequences used for CRISPR mutagenesis**
